## Supplementary Table 1 for "PRR adjuvants restrain high stability peptides presentation on APCs"

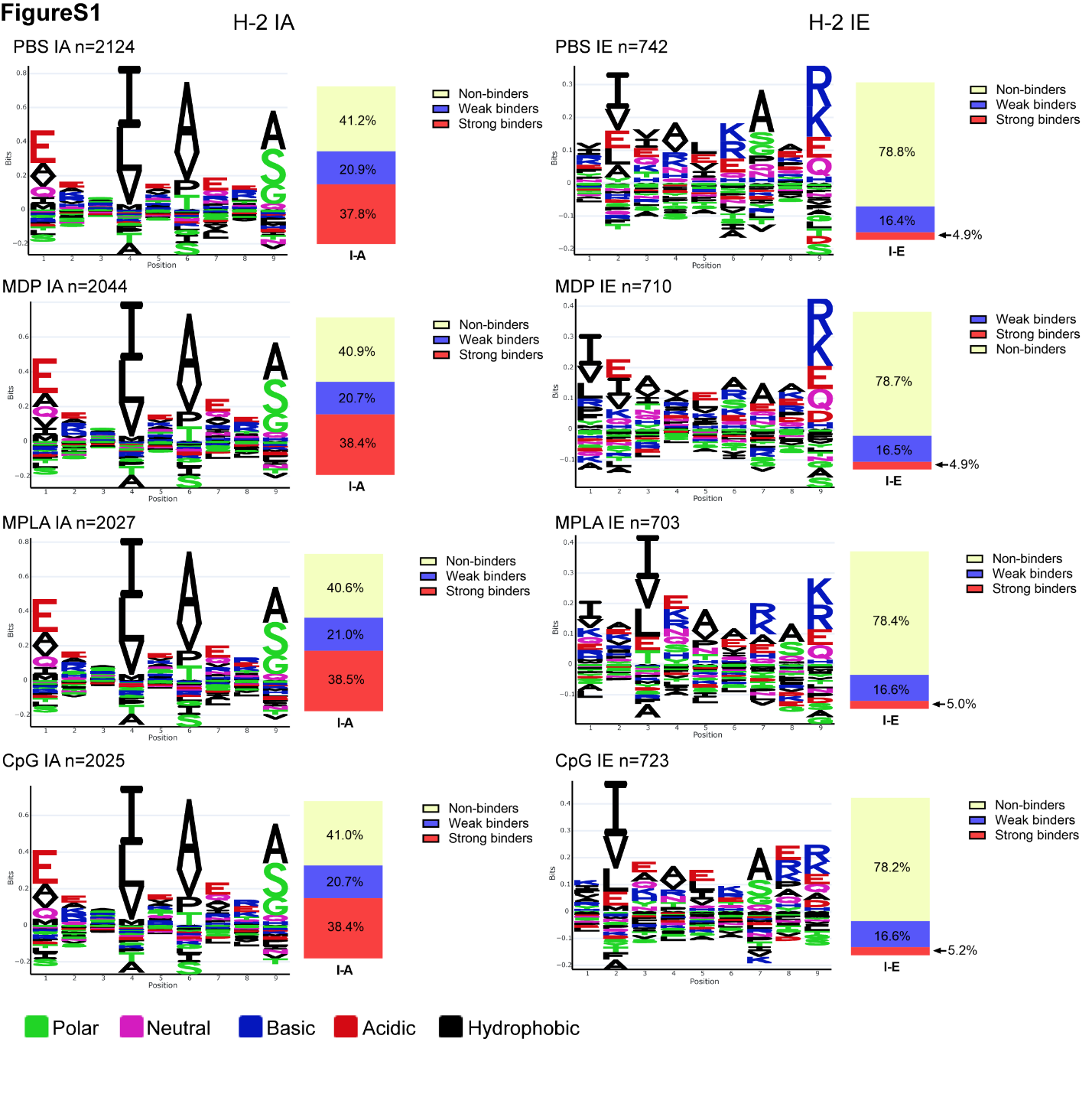

**Fig. S1. Peptide logos and MHC-II binding assigned to individual alleles.**

Logo plots of the identified MHC peptides for individual alleles H2-IA and H2-IE were analyzed. The fractions of MHC peptides between 9–22 mer binding to each allele are shown.

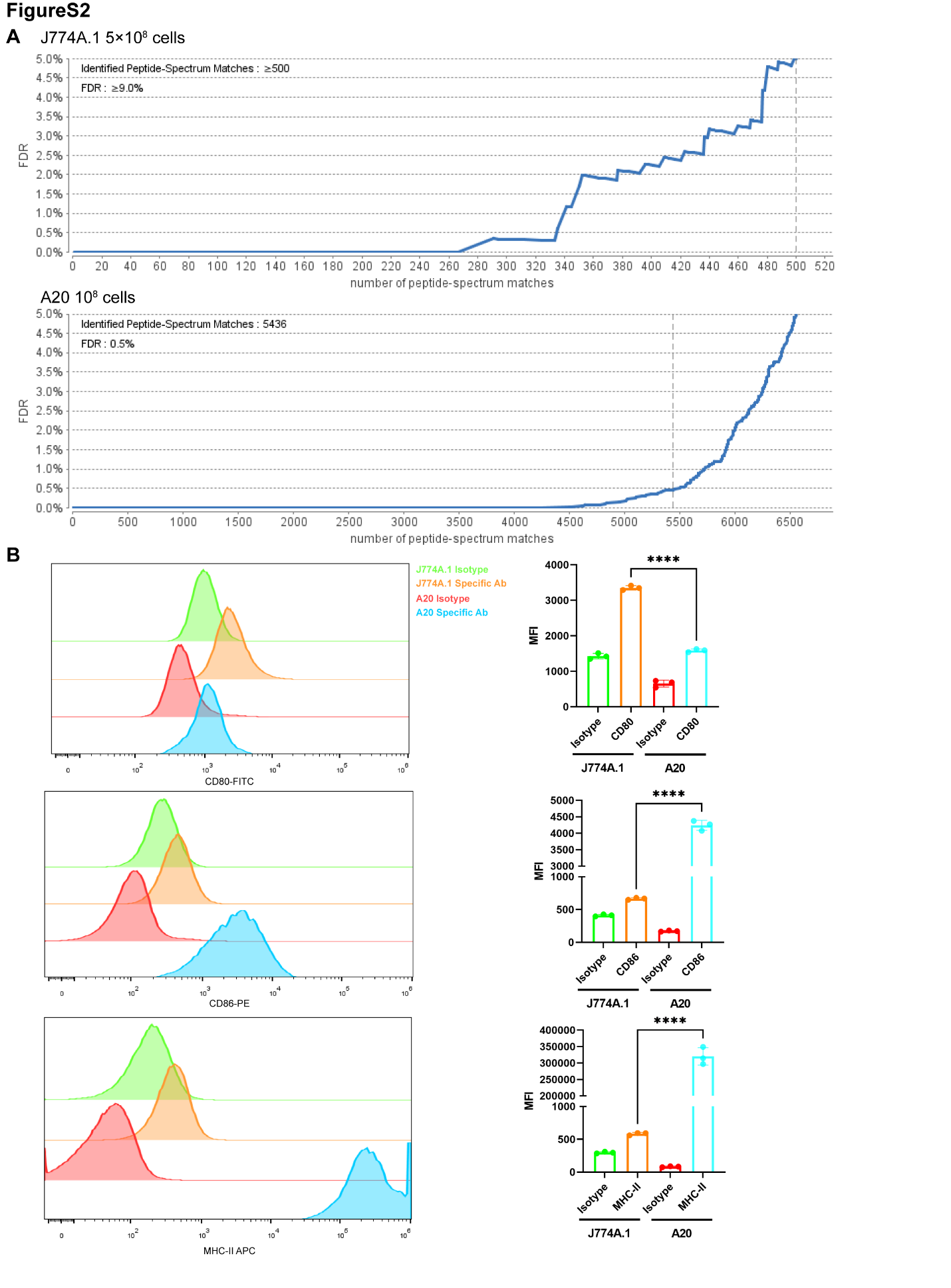

**Fig. S2. Immunopeptidomics of J774A.1 cell line and surface marker detection.**

**(A)** 5×108 J774A.1 cells and 108 A20 cells were used for immunopeptidomics. The number of MHC peptides was compared. Less than 350 MHC peptides in J77A.1 cells and more than 5500 MHC peptides in A20 cells were observed at a peptide spectrum match (PSM) level of <1.0% false discovery rate (FDR). **(B)** J774.1 and A20 cell surface markers were stained with specific antibodies or corresponding isotype controls. Then, expression levels were compared between J774.1 and A20 cells (n=3). MFI, mean fluorescence intensity. *****p*<0.0001.

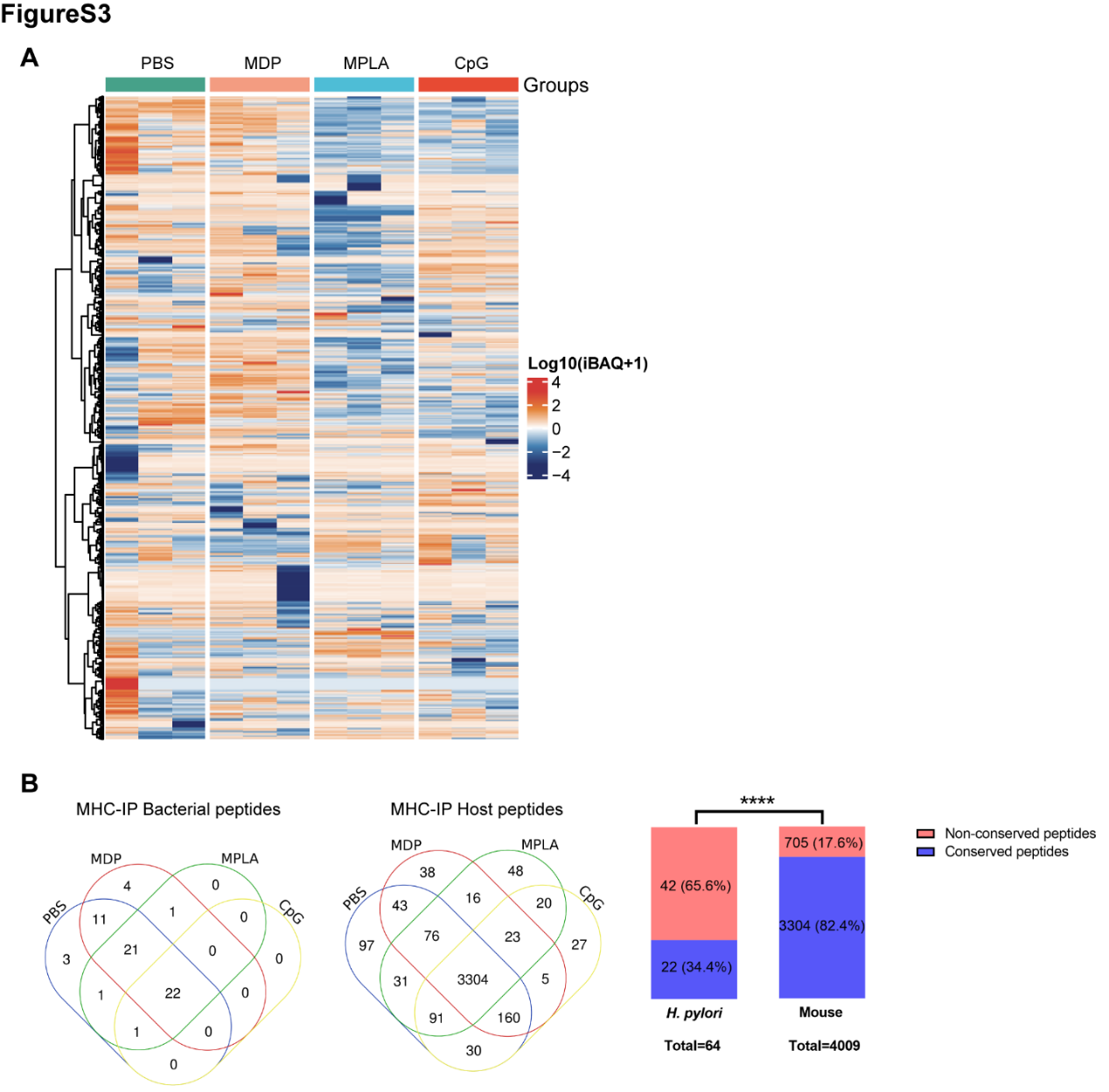

**Fig. S3. Profiling peptides from MHC-II immunopeptidomes.**

**(A)** Heatmap showing the host MHC peptides from the MHC-II immunopeptidome. **(B)** Venn diagrams showing the distribution of bacterial and host MHC peptides in different adjuvant groups. Fractions of bacterial and host conserved and non-conserved MHC peptides were analyzed. The number and percentage of peptides are indicated. *****p*<0.0001.

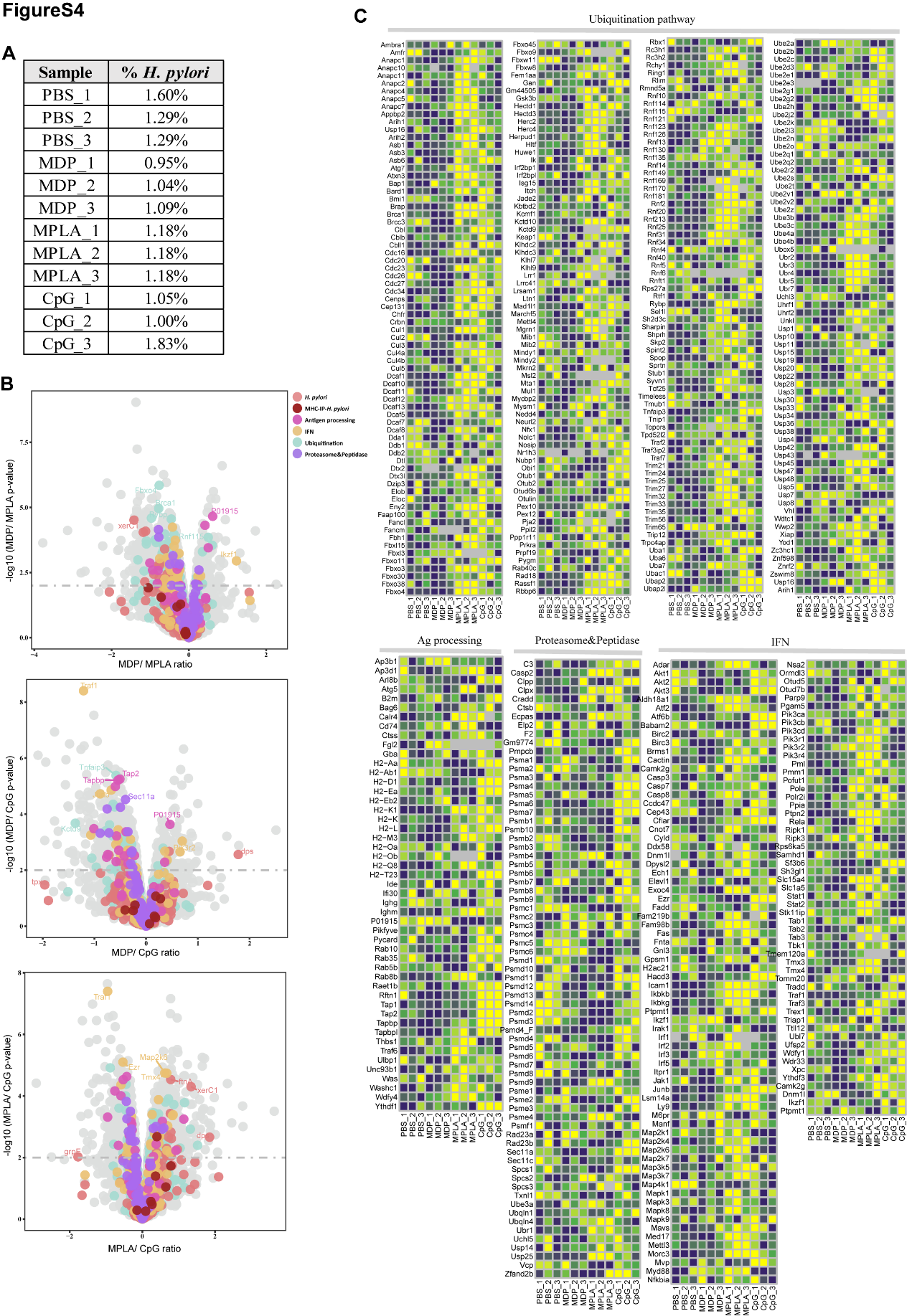

**Fig. S4. Bacterial proteins and host antigen presentation proteins in the whole-proteome of different adjuvant groups.**

**(A)** Percentage of bacterial protein abundances in the total whole proteomes. **(B)** Volcano plots comparing protein levels among adjuvant-treated groups (the dashed line: *p*<0.01). **(C)** Abundances of proteins (log10 protein iBAQ) involved in antigen processing and presentation in the whole-proteome analysis.

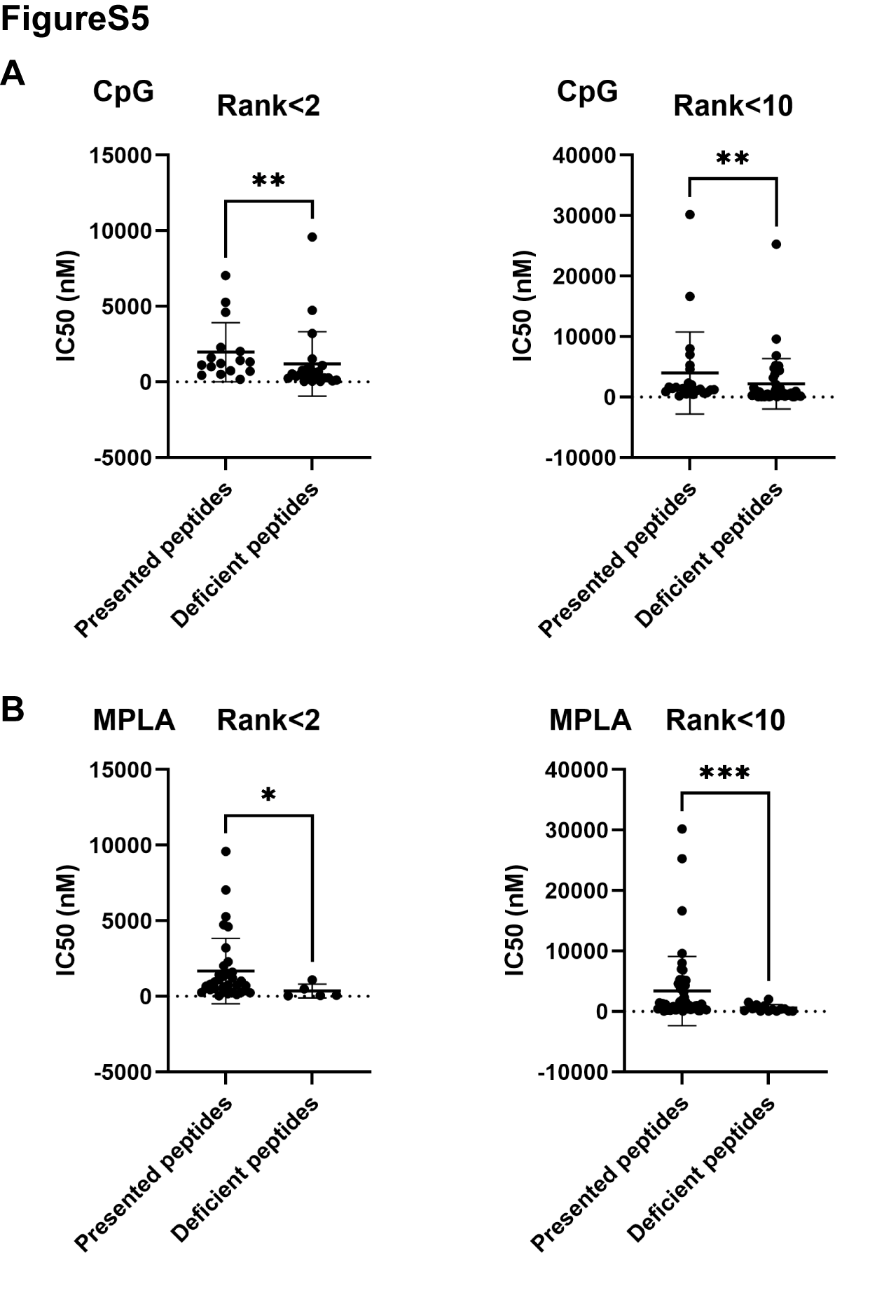

**Fig. S5. Binding affinities of bacterial MHC peptides were assessed at cutoffs of percentile rank <2 (Strong Binders) or <10 (Weak Binders).**

Peptides binding to MHC-II alleles were screened at cutoffs of percentile rank <2 (Strong Binders) or <10 (Weak Binders) by the NetNHCIIpan_el 4.1 method using the IEBD website. The IC50 of these peptides was predicted and compared using the NN alignment method. IC50 of the presented and deficient peptides after **(A)** CpG and **(B)** MPLA stimulation at cutoffs of percentile ranks <2 and <10 were analyzed. **p*<0.05, ***p*<0.01, ****p*<0.001.

Table S1. Key Resources Table

| REAGENT or RESOURCE | SOURCE | IDENTIFIER |
| --- | --- | --- |
| Antibodies |  |  |
| Anti-mouse CD45-FITC | Biolegend | Cat #103108 |
| Anti-mouse CD3-PE-Cy7 | Biolegend | Cat #100320 |
| Anti-mouse IFN-γ- PE/Dazzle594 | Biolegend | Cat #505846 |
| Anti-mouse MHC-II-APC | Biolegend | Cat #107614 |
| Anti-mouse CD4-APC | Biolegend | Cat #100412 |
| Anti-mouse CD86-PE | Biolegend | Cat #105007 |
| Anti-mouse CD80- FITC | Biolegend | Cat #104706 |
| Anti-mouse CD45- APC-Cy7 | Biolegend | Cat #147718 |
| Anti-mouse CD16/32 TruStain FcX™ PLUS | Biolegend | Cat #156604 |
| Anti-Mouse H2-IAd/IEd antibody | BioXcell | Cat #BE00108 |
| Anti- Mouse MHC-II (I-A/I-E) antibody | Thermo Fisher | Cat #14-5321-82 |
| Cells and Strain |  |  |
| A20 cell line | ATCC | Cat #TIB-208 |
| J774A.1 cell line | ATCC | Cat #TIB-67 |
| 11637 *H. pylori* strain | ATCC | Cat #43504 |
| Chemicals, Adjuvants and Peptides |  |  |
| MPLA-SM | Invivogen | Cat #tlrl-mpla2 |
| MDP | Invivogen | Cat #tlrl-mdp |
| CPG ODN | Generay | Customized quote |
| Brefeldin A Solution | Biolegend | Cat #420601 |
| Recombinant Murine IL-2 | PeproTech | Cat #212-12 |
| DMSO | Sigma-Aldrich | Cat #D2650 |
| Fetal calf serum (FBS) | Gibco | Cat #10099141C |
| Fetal Bovine Serum | BI | Cat #04-001-1ACS |
| L-Glutamine | Gibco | Cat #25030081 |
| Aseptic rabbit blood | Sbjbio | Cat #SBJ-ST-RAB002 |
| BASIC PBS, pH 7.4 | Gibco | Cat #C10010500BT |
| T75 cell culture flask | Corning | Cat #430641 |
| CHAPS | Millipore | Cat #1116620001 |
| CNBr-activated Sepharose | Cytivia | Cat #17-0430-01 |
| Glycine | Sigma-Aldrich | Cat #V900144 |
| Tris | Sigma-Aldrich | Cat #V900483 |
| Octyl-β-D-glucopyranaside | Sigma-Aldrich | Cat #V900365 |
| Europium-Streptavidin | Abcam | Cat #ab270228 |
| Hydrochloric acid solution | Xilong Scientific | Cat #7647-01-0 |
| Poly prep chromatography columns | Bio-Rad | Cat #731-1550 |
| Proteases inhibitor | ThermoFisher | Cat #A32959 |
| Sodium Bicarbonate | Macklin | Cat #S799206 |
| Sodium hydroxide | Macklin | Cat #S885179 |
| Glacial acetic acid | Macklin | Cat #A801297 |
| Tris-HCl | Solarbio | Cat #T8230 |
| Water, LC/MS Grade | Thermo Fisher | Cat #W64 |
| Sodium Hydroxide Solution | Macklin | Cat #S885179 |
| Acetonitrile，LC/MS Grade LC-MS | Aladdin | Cat #A120771 |
| Methanol，LC/MS Grade | Aladdin | Cat #M116125 |
| Ethanol absolute | Sinopharm | Cat #10009218 |
| Sodium Chloride | Sinopharm | Cat #10019318 |
| Isopropanol | Sinopharm | Cat #80109218 |
| 2-Mercaptoethanol (1000×), liquid | Gibco | Cat #21985 |
| Penicillin-Streptomycin, liquid | Gibco | Cat #15140122 |
| DMEM | Gibco | Cat #C11995500BT |
| RPMI 1640 | Gibco | Cat #C11875500BT |
| RIPA lysis and extraction buffer | Thermo Fisher | Cat #89900 |
| Halt™ Protease inhibitor mixture (100×) | Thermo Fisher | Cat #78430 |
| PMSF Protease inhibitor | Thermo Fisher | Cat #36978 |
| Citric acid | MCE | Cat #HY-N1428 |
| Mouse 1× Lymphocyte Separation Medium | Dakewe | Cat #7211011 |
| Complete Freund's Adjuvant, CFA | Sigma-Aldrich | Cat #F5881-10ML |
| Mouse IFN-γ precoated ELISpot kit | Dakewe | Cat #2210003 |
| Ovalbumin (OVA) | Yuanye Biotech | Cat #9006-59-1 |
| PMA | BioGems | Cat #1652981 |
| Purified mouse MHC-II | Proimmune | Customized quote |
| Synthetic peptides, > 90% purity | China Peptides | Customized quote |
| Critical commercial assays |  |  |
| MHC-II peptidome LC-MS/MS | Wininnovate Bio | Customized quote |
| Whole proteome LC-MS/MS | Wininnovate Bio/ PTM Bio | Customized quote |
| Software and algorithms |  |  |
| MHC-II binding prediction | IEDB | https://www.iedb.org/ |
| Prediction of peptide-MHC-II binding motifs | MhcVizPipe (MVP) | https://github.com/CaronLab/MhcVizPipe/ |

Table S2. Whole-proteome data, related to Figure 5 and Figure S4.

The whole proteome data of A20 cells treated with different adjuvants at 12h. The “Gene anno” sheet contains the KEGG and GO annotations of proteins from antigen processing, peptidase function, ubiquitination pathway, interferon (IFN) signaling and *H. pylori*. The “iBAQ” sheet contains the iBAQ values of each protein in different adjuvant groups.
